# Systems biology analysis of mitogen activated protein kinase inhibitor resistance in malignant melanoma

**DOI:** 10.1101/231142

**Authors:** Helma Zecena, Daniel Tveit, Zi Wang, Ahmed Farhat, Parvita Panchal, Jing Liu, Simar J. Singh, Amandeep Sanghera, Ajay Bainiwal, Shuan Y. Teo, Frank L Meyskens, Feng Liu-Smith, Fabian V. Filipp

## Abstract

Kinase inhibition in the mitogen activated protein kinase (MAPK) pathway is a standard therapy for cancer patients with activating BRAF mutations. However, the anti-tumorigenic effect and clinical benefit are only transient, and tumors are prone to treatment resistance and relapse. To elucidate mechanistic insights into drug resistance, we have established an *in vitro* cellular model of MAPK inhibitor resistance in malignant melanoma. The cellular model evolved in response to clinical dosage of BRAF inhibitor, vemurafenib, PLX4032. We conducted transcriptomic expression profiling using RNA-Seq and RT-qPCR arrays. Pathways of melanogenesis, MAPK signaling, cell cycle, and metabolism were significantly enriched among the set of differentially expressed genes of vemurafenib-resistant cells vs control. The underlying mechanism of treatment resistance and pathway rewiring based on non-genomic adaptation was validated in two distinct melanoma models, SK-MEL-28 and A375. Both cell lines have activating BRAF mutations and display metastatic potential. Downregulation of tumor suppressors and negative MAPK regulators, dual specific phosphatases, reengages mitogenic signaling. Upregulation of growth factors or cytokine receptors triggers signaling pathways circumventing BRAF blockage. Changes in amino acid and one-carbon metabolism support cellular proliferation despite MAPK inhibitor treatment. In addition, an upregulation of pigmentation in inhibitor resistant melanoma cells was observed. Cellular pathways utilized during inhibitor resistance promoted melanogenesis, a pathway which partially overlaps with MAPK signaling. Upstream regulator analysis suggested gene expression changes of forkhead box and hypoxia inducible factor family transcription factors. The established cellular models offer mechanistic insight into cellular changes and therapeutic targets under inhibitor resistance in malignant melanoma. At a systems biology level, the MAPK pathway undergoes major rewiring while acquiring inhibitor resistance. The outcome of this transcriptional plasticity is selection for a set of transcriptional master regulators, which circumvent upstream targeted kinases and provide alternative routes of mitogenic activation. A fine-woven network of redundant signals maintains similar effector genes allowing for tumor cell survival and malignant progression in therapy resistant cancer.

## Introduction

Cancer drug resistance is a major obstacle to achieve durable clinical responses with targeted therapies. This highlights a need to elucidate the underlying mechanisms responsible for resistance and identify strategies to overcome this challenge. In malignant melanoma, activating point-mutations in the mitogen activated protein kinase (MAPK) pathway in BRAF kinase (B-Raf proto-oncogene, serine/threonine kinase, Gene ID: 673) [1-3] made it possible to develop potent kinase inhibitors matched to genotyped kinase mutations in precision medicine approaches [4-6]. In tumors expressing the oncoprotein BRAF(V600E), the inhibitor molecules vemurafenib, dabrafenib, and encorafenib are designed to lock the ATP binding site into an inactive conformation of the kinase [4], the preferred state of wild-type RAF proteins. Trametinib and cobimetinib are targeting MAP2K7 (MEK, mitogen-activated protein kinase kinase 7, Gene ID: 5609), the BRAF target and downstream effector molecule. Combinations of single BRAF inhibitors (BRAFi) have proved to be superior to single-agent regimens [7-10]. For both groups, combinations proved to be superior to single-agent regimens: BRAF inhibitors (BRAFi) in combination with MEK inhibitors (MEKi) improved survival compared to single MAPK inhibitors (MAPKi). However, many patients responding to small molecule inhibition of the MAPK pathway will develop resistance. Ultimately, disease progression will take place and patients relapse with lethal drug-resistant disease.

Acquired resistance has been shown to involve a diverse spectrum of oncogenic mutations in the MAPK pathway [11-15]. In addition, non-genomic activation of parallel signaling pathways was noted [16]. Cell-to-cell variability in BRAF(V600E) melanomas generates drug-tolerant subpopulations. Selection of genetically distinct, fully drug-resistant clones arise within a set of heterogeneous tumor cells surviving the initial phases of therapy due to drug adaptation [17]. Non-genomic drug adaptation can be accomplished reproducibly in cultured cells, and combination therapies that block adaptive mechanisms *in vitro* have shown promise in improving rates and durability of response [18]. Thus, better understanding of mechanisms involved in drug adaptation is likely to improve the effectiveness of melanoma therapy by delaying or controlling acquired resistance.

## Methods

### Cellular models of malignant melanoma

SK-MEL-28 and A375 are human skin malignant melanoma cell lines with BRAF(V600E) activation that are tumorigenic in xenografts [19-22] (HTB-72 and CRL-1619, American Type Culture Collection, Manassas, VA). The cell lines are maintained in DMEM medium supplemented with 10% fetal bovine serum and antibiotics (10-017-CV, 35-010-CV, 30-002-CI Corning, Corning, NY). All experimental protocols were approved by the Institutional Review Boards at the University of California Merced and Irvine. The study was carried out as part of IRB UCM13-0025 of the University of California Merced and as part of dbGap ID 5094 on somatic mutations in cancer and conducted in accordance with the Helsinki Declaration of 1975.

BRAFi-resistant (BRAFi-R) models were obtained by challenging cancer cell lines with incrementally increasing vemurafenib (PLX4032, PubChem CID: 42611257, Selleckchem, Houston, TX) concentrations in the culture media. Starting at 0.25 μM, which matched the naïve half maximal inhibitory concentration (IC50) of the parental cell lines, the vemurafenib concentrations were increased every 7 days in an exponential series up to 100-fold the naïve IC50 concentrations. Following this 6-week selection protocol, vemurafenib-adapted, BRAFi-resistant models were maintained in media supplemented with 5.0 μM vemurafenib.

### Transcriptomic profiling and differential gene expression analysis

Total RNA from malignant melanoma cells was extracted using a mammalian RNA mini preparation kit (RTN10-1KT, GenElute, Sigma EMD Millipore, Darmstadt, Germany) and then digested with deoxyribonuclease I (AMPD1-1KT, Sigma EMD Millipore, Darmstadt, Germany). Complementary DNA (cDNA) was synthesized using random hexamers (cDNA SuperMix, 95048-500, Quanta Biosciences, Beverly, MA). The purified DNA library was sequenced using a HighSeq2500 (Illumina, San Diego, CA) at the University of California Irvine Genomics High-Throughput Facility. Purity and integrity of the nucleic acid samples were quantified using a Bioanalyzer (2100 Bioanalyzer, Agilent, Santa Clara, CA). Libraries for next generation mRNA transcriptome sequencing (RNA-Seq) analysis were generated using the TruSeq kit (Truseq RNA Library Prep Kit v2, RS-122-2001, Illumina, San Diego, CA). In brief, the workflow involves purifying the poly-A containing mRNA molecules using oligo-dT attached magnetic beads. Following purification, the mRNA is chemically fragmented into small pieces using divalent cations under elevated temperature. The cleaved RNA fragments are copied into first strand cDNA using reverse transcriptase and random primers. Second strand cDNA synthesis follows, using DNA polymerase I and RNase H. The cDNA fragments are end repaired by adenylation of the 3’ ends and ligated to barcoded adapters. The products are then purified and enriched by nine cycles of PCR to create the final cDNA library subjected to sequencing. The resulting libraries were validated by qPCR and size-quantified by a DNA high sensitivity chip (Bioanalyzer, 5067-4626, Agilent, Santa Clara, CA). Sequencing was performed using 50 base pair read length, single-end reads, and more than 10^7^ reads per sample. Raw sequence reads in the file format for sequences with quality scores (FASTQ) were mapped to human reference Genome Reference Consortium GRCh38 using Bowtie alignment with an extended Burrows-Wheeler indexing for an ultrafast memory efficient alignment [23]. Read counts are scaled via the median of the geometric means of fragment counts across all libraries. Transcript abundance was quantified using normalized single-end RNA-Seq reads in read counts or reads per kilobase million (RPKM). Since single-end reads were acquired in the sequencing protocol, quantification of reads or fragments yields identical results. Statistical testing for differential expression was based on read counts and performed using EdgeR in the Bioconductor toolbox [24]. Differentially expressed genes were further analyzed using Ingenuity Pathways Analysis (IPA, Qiagen, Rewood City, CA), classification of transcription factors (TFClass) and gene set enrichment analysis (GSEA, Broad Institute, Cambridge, MA) [25, 26]. Triple replicate samples were subjected to SYBR green (SYBR green master mix, PerfeCTa^®^ SYBR^®^ Green FastMix^®^, 95072-05k, Quanta Biosciences, Beverly, MA) real time quantitative polymerase chain reaction (RT-qPCR) analysis in an Eco system (Illumina, San Diego). RT-qPCR threshold cycle (CT) values were normalized using multiple housekeeping genes like actin beta (ACTB, GeneBank: 60), cyclophilin A (PPIA, peptidylprolyl isomerase A, GeneBank: 5478) and RNA polymerase II subunit A (POLR2A, GeneBank: 5430). Gene expression profiles were analyzed using the ∆∆CT method. Oligo nucleotides crossing exon-exon-junctions of transcripts served for RT-qPCR validation of RNA-Seq signals of differentially expressed target genes in BRAFi-resistant melanoma cells (Supplementary table 1).

### Inhibitor cytotoxicity studies

Chemical BRAFi against BRAF(V600E), vemurafenib, was dissolved in dimethyl sulfoxide (DMSO, Sigma) as 10.0 mM stock solution and used in treatments in final concentrations between 0.01 μM and 50.0 μM. Melanoma control experiments were carried out in the presence of equivalent amounts of DMSO solvent without drug. Cell viability was determined using a 3-(4,5-dimethylthiazol-2-yl)-2,5-diphenyltetrazolium bromide (MTT, M6494, Thermo Fisher Scientific, Waltham, MA) absorbance assay by subtracting background readout at 650 nm from response readout at 570 nm wavelength. IC50 concentrations were determined after 72 hours of drug treatment between 0.01-100 μM in two-fold dilution series. Analysis was performed using CalcuSyn (2.0, Biosoft, Cambridge, UK).

### Melanin quantification

Melanin pigment production of cultured cells was determined by colorimetric measurements normalized for total protein levels in arbitrary units [27, 28]. Melanoma cells were harvested by centrifugation at 3,000 rpm (3830 g, Z326K, Labnet International, Edison, NJ) and dissolved in either 1.0 N NaOH for melanin assay or lysis 250 for protein assay. The cell lysates were sonicated, incubated at room temperature for 24 hours, and cleared by centrifugation at 13,000 rpm for 10 minutes (17,000 g, Z326K, Labnet International, Edison, NJ). The absorption of the supernatant was measured at OD475 in a spectrophotometer (Smartspec3000, Bio-Rad, Carlsbad, CA). Cells were lysed in mild denaturing conditions in lysis 250 buffer (25 mM Tris, [pH 7.5], 5 mM EDTA, 0.1% NP-40, 250 mM NaCl) containing proteinase inhibitors (10 μg/ml aprotinin, 10 μg/ml leupeptin, 10 μg/ml pepstatin, 5 μg/ml antipain, 1 mM phenylmethylsulfonyl fluoride). The total protein amount in the lysates was quantified using a colorimetric Bradford assay (5000001, Bio-Rad, Richmond, CA) at 595 nm and an incubation time of 30 min [29].

## Results

### Generation of BRAFi-resistant melanoma cell lines

The parental melanoma cell lines SK-MEL-28 and A375 were exposed to incrementally increasing concentrations of the mutant-BRAF inhibitor vemurafenib (Figure 1A). At the initial inhibitor concentration matching the IC50 of vemurafenib in the naïve parental melanoma cells [30, 11] cell proliferation slowed down. Surviving cells were propagated and subjected to an exponential series of increasing vemurafenib concentrations until BRAFi-R sublines were obtained tolerating at least 5 μM vemurafenib in the culture media with similar cell proliferation rates as the parental cell lines of 0.67 doublings per day.

**Figure 1:**
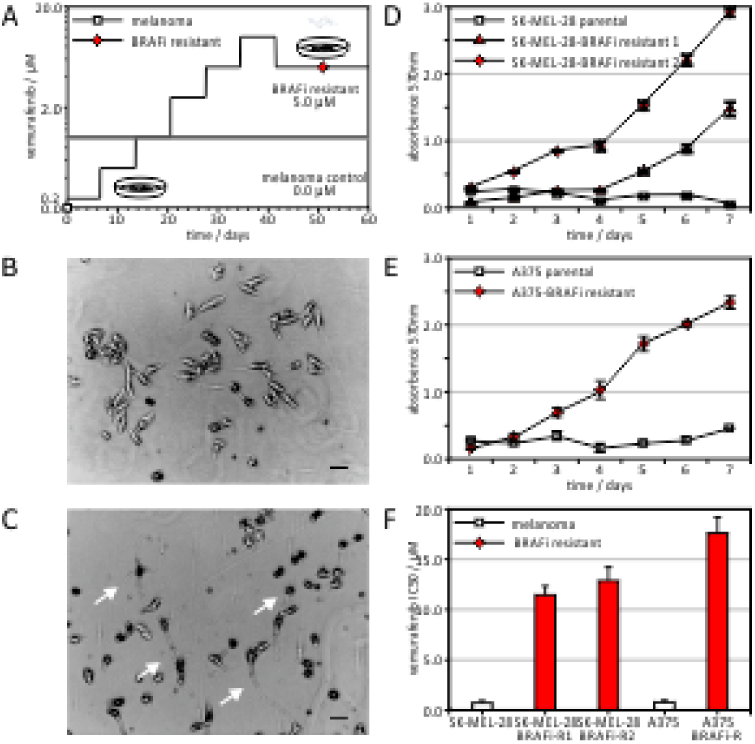
Establishing mitogen activated protein kinase inhibitor resistant melanoma models

Some BRAFi-R cell lines showed structures typically observed in differentiated melanocytes (Figure 1B-C). In the presence of 5 μM vemurafenib, however, the parental cells were not able to grow but the resistant cells proliferated comparable to naïve cell lines (Figure 1D-E). For the SK-MEL-28 cell line, two resistant sublines were established. The resistant sublines displayed IC50 values of 11.5 ± 0.9 μM and 13.3 ± 1.2 μM for SK-MEL-28-BRAFi-R1 and SK-MEL-28-BRAFi-R2 respectively, which is approximately 10-20 fold of the IC50 in a low micro-molar range for the parental cells with 0.74 ± 0.05 μM. For the A375 cell line, the IC50 of the A375-BRAFi-R cell line was observed at 17.7 ± 1.5 μM, 22.7 fold of IC50 for the parental A375 cells with 0.78 ± 0.22 μM (Figure 1F).

### Transcriptomic profiling identifies non-genomic rewiring of treatment-resistant cancer cells

We conducted transcriptomic gene expression profiling of BRAFi treatment-resistant SK-MEL-28-BRAFi-R1 and SK-MEL-28-BRAFi-R2 cell lines by RNA-Seq and looked for differential expression versus the parental SK-MEL-28 cell line. In total, 980 unique transcripts showed significant differential expression in RNA-Seq experiments with p-values below 0.05, absolute log-fold change (LOG(FC)) greater or equal 1.0 (Figure 2A-B). The differentially expressed genes included 505 upregulated transcripts and 475 downregulated transcripts (Supplementary table 2-3). We subjected the identified directional sets to pathway enrichment analysis (Supplementary table 4). Distinct clusters stood out and showed significant enrichment with p-values below 0.05 and q-values below 0.10 (Figure 2C). Melanogenesis and pathways in cancer, inflammation, nuclear factor kappa-light-chain-enhancer of activated B cells (NFκB) and signal transducer and activator of transcription (STAT) signaling, metabolic pathways including alanine, tyrosine, valine, leucine, inositol, one-carbon metabolism, cell-adhesion molecules, neurotrophin signaling were over-represented in the upregulated dataset. MAPK signaling and epithelial-mesenchymal transition (EMT) were differentially expressed and characterized by both strong up- and downregulation. Extra-cellular matrix (ECM) receptors, cell cycle, and hypoxia signaling were enriched in the downregulated dataset. Of the 980 differential expressed genes, we validated expression changes of 150 genes by RT-qPCR (Figure 2D, Supplementary table 3). A majority of 64.0% (96 of 150) of the transcript regulation in treatment-resistant melanoma in the RT-qPCR data corresponded significantly in direction with the RNA-Seq data upon treatment resistance with p-values below 0.05. When both treatment resistance models of SK-MEL-28 and A375 were taken into consideration, about half of the tested genes, 50 of 96, showed consistent regulation (Figure 2E, Supplementary table 3). Genes in MAPK signaling included nuclear factor of activated T-cells 2 (NFATC2, Gene ID: 4773), phospholipase A2 group VI (PLA2G6, Gene ID: 8398), dual specificity phosphatase 1 (DUSP1, Gene ID: 1843), and dual specificity phosphatase 2 (DUSP2, Gene ID: 1844), which were downregulated in the BRAFi-resistant cells compared to control. Genes contributing to melanogenesis adenylate cyclase 1, (ADCY1, Gene ID: 107), dopachrome tautomerase (DCT, TYRP2, Gene ID: 1638), and platelet derived growth factor C (PDGFC, Gene ID: 56034) were upregulated. Lastly, metabolic regulators such as methylenetetrahydrofolate dehydrogenase 2, (MTHFD2, Gene ID: 10797) for folate metabolism, asparagine synthetase (ASNS, Gene ID: 440) for amino acid metabolism, and NME/NM23 nucleoside diphosphate kinase 1 (NME1, Gene ID: 4830) and dihydropyrimidine dehydrogenase (DPYD, Gene ID: 1806) for pyrimidine metabolism were significantly upregulated (Figure 2D). Taken together, the adaptive transcriptomic changes were validated in two distinct melanoma models, SK-MEL-28 and A375, both cell lines with metastatic potential showed differential expression of MAPK signaling while activating alternative receptor interactions and metabolic processes.

**Figure 2:**
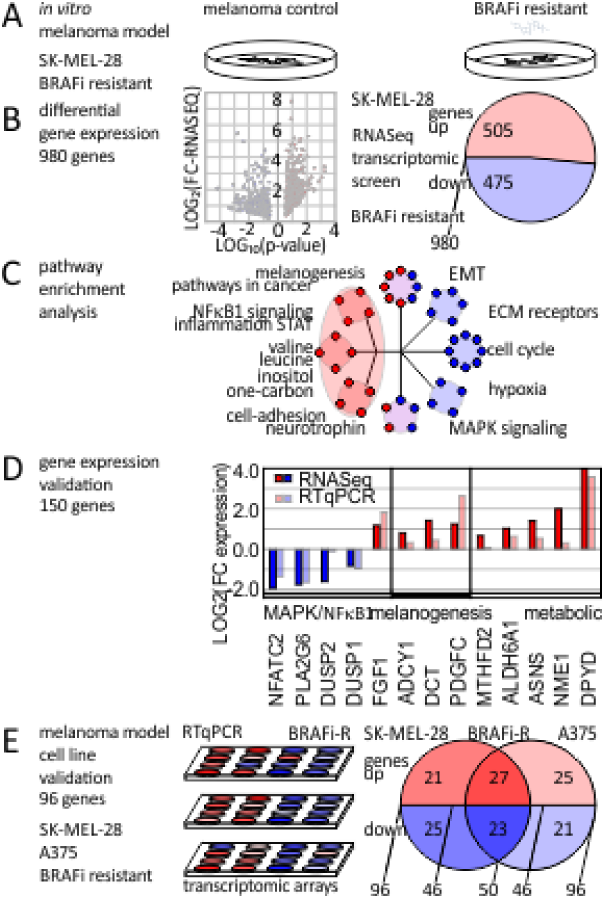
Transcriptomic profiling of BRAF inhibitor resistance in cellular models of malignant melanoma

### Upstream regulator analysis suggests control by transcription factor families

In a next step, we subjected the gene list to hierarchical transcription factor motif analysis. We asked whether any of the enriched transcription factor motif families were represented in the differential gene expression data. In detail, we looked for transcription factors as well as their target genes whose promoters show the respective transcription factor binding sites are among the same list of regulated genes (Figure 3A). It is expected that differentially expressed transcription factors show enrichment in significantly deregulated target genes. Further, identified target genes with enriched transcription factor motifs will have major contributions to significantly deregulated pathways under treatment resistance (Figure 3B). A network illustration of transcriptional master regulators, target genes, and dysregulated effector network upon treatment resistance demonstrates transcriptional synergy (Figure 3C). Upregulated transcription factor families included Rel homology region (RHR) NFκB-related factors, forkhead box (FOX), Zinc finger E-box-binding homeobox domain factors (ZEB), nuclear steroid hormone receptor subfamily 3 (NR3C, androgen receptor and progesterone receptor), hypoxia-inducible and endothelial PAS domain-containing factors (HIF, EPAS), and the cell cycle transcription factor family (E2F) (Figure 3B). Downstream enriched target genes comprised members of interleukin (IL), chemokine receptor (CXCL), matrix metallo proteinase (MMP) families, transcription factors forkhead box O1 (FOXO1, Gene ID: 2308), endothelial PAS domain protein 1 (EPAS1, HIF2A, Gene ID: 2034) and melanogenesis associated metabolic genes, tyrosinase (TYR, OCA1, Gene ID: 7299), DCT, and melanosomal transmembrane protein (OCA2, oculocutaneous albinism II, Gene ID: 4948). Downregulated transcription factors included forkhead box F2 (FOXF2, Gene ID: 2295), which has DUSP2 or transforming growth factor beta 3 (TGFB3, Gene ID: 7043) as target genes. Upstream regulator analysis suggested gene expression changes of nuclear factor kappa B subunit 1 (NFKB1, Gene ID: 4790, V$NFKB_Q6, motif M11921) in complex with REL proto-oncogene (REL Gene ID: 5966, V$CREL_01, motif M10143), EMT modulator zinc finger E-box binding homeobox 1 (ZEB1, Gene ID: 6935, V$AREB6_01, M11244), forkhead box (V$FOXO1_01, motif M11512), and hypoxia inducible factor family transcription factors (V$HIF1_Q3, motif M14011) as master regulators of transcriptional effector networks upon BRAFi treatment resistance.

**Figure 3:**
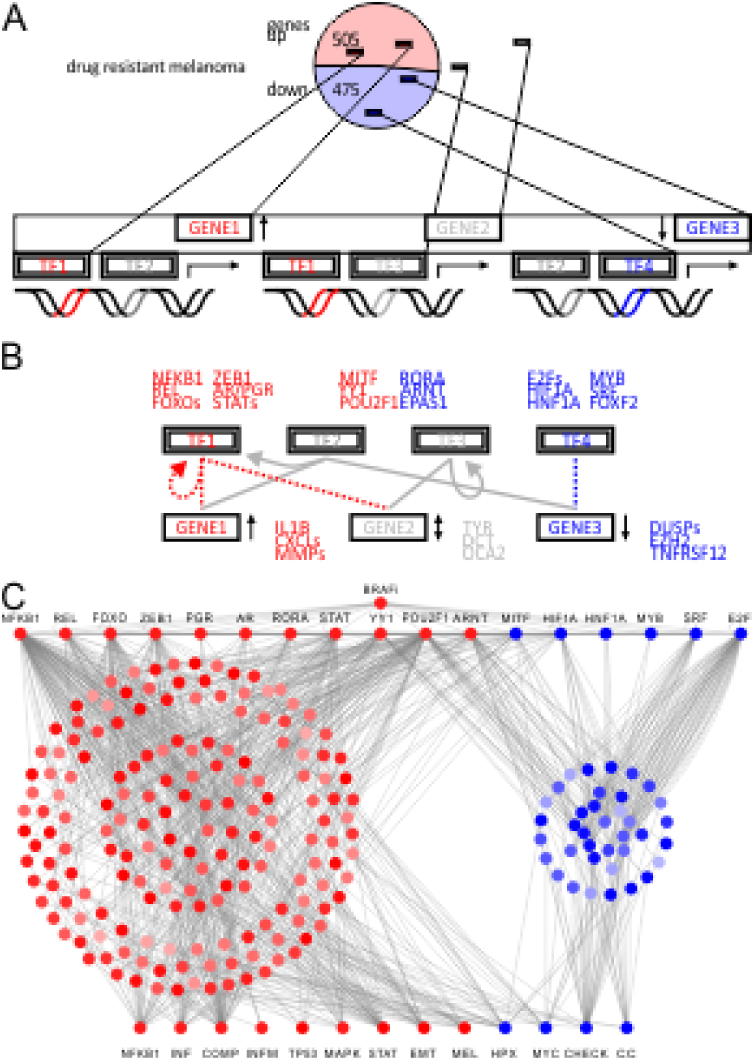
Transcription factor motif analysis of mitogen activated protein kinase inhibitor resistance in cellular models of malignant melanoma

### Validation of pathway rewiring in drug resistance in multiple cell lines by transcriptomics arrays

Transcriptome analysis of reversible drug resistance identified distinct pathways that allowed for circumvention of BRAF blockage (Figure 4A). Cell-to-cell variability in combination with drug exposure selects for distinct sub-populations of MAPKi resistant cell lines. In a hierarchical fashion, transcriptional master regulators promote a distinct set of target genes resulting in circumvention of MAPK inhibition. Receptor activation by fibroblast growth factor 1 (FGF1, Gene ID: 2246) or platelet derived growth factor C (PDGFC, Gene ID: 56034) can lead to activated receptor tyrosine signaling parallel to canonical MAPK signaling [16] (Figure 4B). In addition, downregulation of tumor suppressors reengages mitogenic signaling. The dual specific phosphatases, DUSP1 and DUSP2, have the ability to switch MAPK signaling off and rank among the top downregulated hits. Thus, downregulation of dual specific phosphatases facilitates and reinforces alternative MAPK effector activation under BRAF blockage (Figure 4B). One of the mitogen-activated protein kinase 1 (MAPK1, ERK2, Gene ID: 5594) effector targets, transcription factor EPAS1, showed upregulation and the ability to maintain its transcriptional program. Pro-apoptotic program of TGFB3 was downregulated and included SMAD family member 9 (SMAD9, Gene ID: 4093) and DUSP1/2 (Figure 4C). Adenylate cyclase, G-protein, and phospholipase signaling are alternative cascades observed in cutaneous and uveal melanoma (Figure 4D). Upregulation of ADCY1, endothelin receptor type B (EDNRB, Gene ID: 1910), PLCB4, and cAMP responsive element binding protein 3 (CREB3, Gene ID: 10488) promote MITF activity, the master transcription factor for pigmentation genes. Downstream metabolic enzymes, TYR and DCT, are both MITF target genes and contribute to enhanced eumelanin production observed in some MAPKi resistant cells. The observed pigmentation showed a wide range of from 1.3-fold to up to 16.8-fold upregulation (Figure 4D). While both cell lines showed dysregulation of melanogenesis, involved regulators and effectors were different. SK-MEL-28-BRAFi-R2 has ASIP prominently expressed (TYR (2.1), DCT (2.8), tyrosinase related protein 1 (TYRP1, OCA3, Gene ID: 7306) (0.5), MITF (0.7), agouti signaling protein (ASIP, Gene ID: 434) (18.9)), while A375-BRAFi-R showed strongest regulation of TYRP1 and MITF (TYR (0.34), DCT (0.24), TYRP1 (41.8), MITF (2.94), ASIP (0.41)). In summary, upregulation of growth factors or receptors triggers signaling pathways circumventing BRAF blockage. Changes in amino acid and one-carbon metabolism support cellular proliferation despite inhibitor treatment. In addition, alternative MAPK signaling coincides with differential response of melanogenesis and pigmentation pathways, which partially overlap with MAPK effectors. In particular, NFKB1, REL, ZEB1, FOXO1, and EPAS1 may serve as master regulators to concert broad transcriptional changes implemented in altered cascades of MAPK, TGFB, ADCY, and MITF signaling.

**Figure 4:**
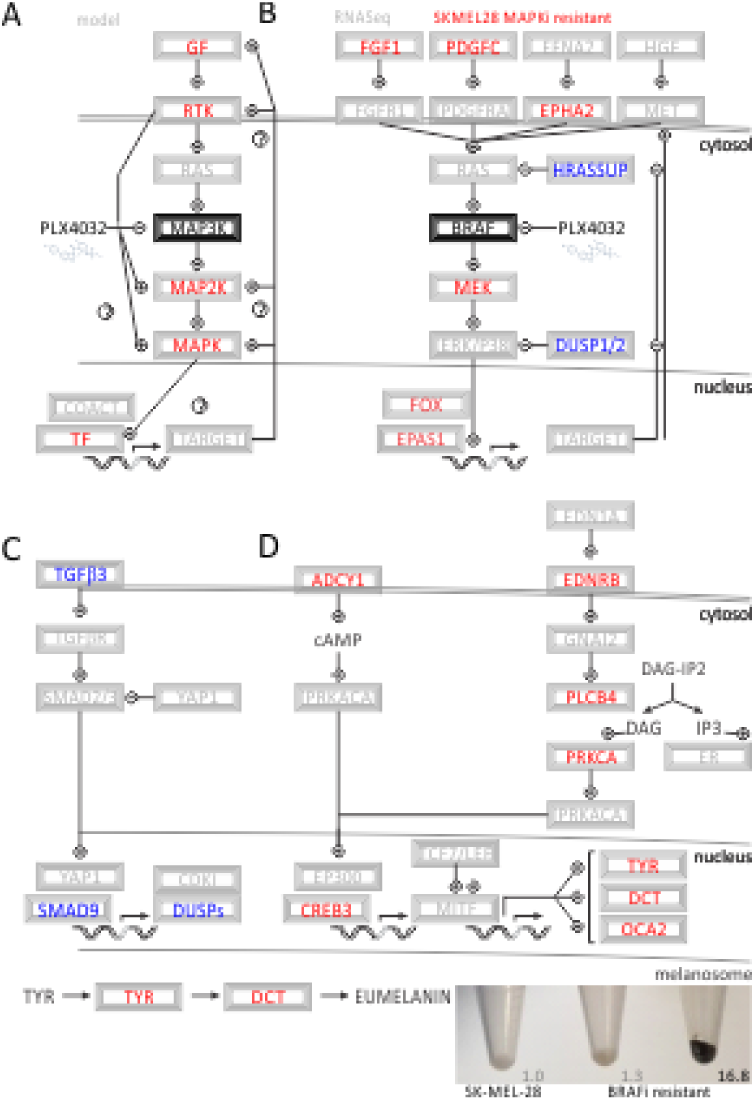
Pathway analysis of BRAF kinase inhibitor resistance shows alternative activation of MAPK targets and pigmentation

## Discussion

Activation of the MAPK pathway is the central and most common oncogenic event in the pathogenesis of malignant melanoma [3, 31]. About 50% of all melanoma patients have activating somatic mutations in the activator loop involving L597, T599, V600, and K601 switching proto-oncogene BRAF into a constitutively active protein kinase and cancer driver. Such activation is supported by somatic copy number amplifications of chromosome 7 [32], often coinciding with somatic V600E/G/K/M/R mutations. Another 20-30% of the patients show non-genomic activation of BRAF by transcriptional upregulation or post-translational modification induced by somatic mutations of upstream signaling molecules like KIT proto-oncogene receptor tyrosine kinase (KIT, Gene ID: 3815), neuroblastoma RAS viral oncogene homolog (NRAS, Gene ID: 4893), or loss-of-function neurofibromin 1 (NF1, Gene ID: 4763). Constitutively activated BRAF phosphorylates MAPK1 and downstream kinases resulting in mitogenic signaling, proliferation, and cell growth. Integrated into this cellular program is negative feedback resulting in reduction of NRAS expression [33, 34].

Genomic sequencing has facilitated the understanding of acquired resistance mechanisms to MAPKis [14, 35, 36, 15, 37, 16, 38]. Detected genetic aberrations included mutations in NRAS, MAPK1/2, phosphatidylinositol-4,5-bisphosphate 3-kinase catalytic subunit alpha (PIK3CA, Gene ID: 5290), and phosphatase and tensin homolog (PTEN, Gene ID: 5728). Somatic melanoma mutations provide examples of how single, well-defined genomic events can confer resistance against vemurafenib treatment. In contrast, transcriptomic as well as epigenomic regulation can provide insight into resistance states that may involve larger networks. Eventually, neither genomic nor transcriptomic readouts will be supported by kinome data reporting on resistance states of phosphoproteins that can be modulated by post-translational modification.

The transcriptomic profiles revealed a network of genes involved in adenylate cyclase signaling conferring resistance and contributing to melanogenesis. ADCY1 and CREB3 are prominent members of the melanogenesis pathway exhibiting mitogenic control and MITF activation. Similarly, a gain-of-function screen confirmed a cyclic-AMP-dependent melanocytic signaling network including G-protein-coupled receptors, adenylate cyclase, protein kinase cAMP-activated catalytic subunit alpha (PRKACA, Gene ID: 5566), and cAMP responsive element binding protein 1 (CREB1, Gene ID: 1385) [39]. The MAPK pathway negatively regulates MITF protein level as well as activity [28], which in turn regulates a series of cell cycle regulating genes. In particular, P16INK4A and P21CIP1, gene products of cyclin dependent kinase inhibitor 2A (CDKN2A, Gene ID: 1029) and cyclin dependent kinase inhibitor 1A (CDKN1A, Gene ID: 1026), respectively, differentiation genes TYR, DCT, TYRP1 as well as survival genes B-cell lymphoma 2 apoptosis regulator (BCL2, Gene ID: 596) and BCL2 family apoptosis regulator (MCL1, Gene ID: 4170) are effector genes under the control of MITF. Inhibition of MITF resulted in sensitivity to chemotherapy drugs [40]. In contrast, upregulation of MITF in therapy-resistance may present itself as a survival mechanism, which coincides with upregulation of melanin, hence it may serve as prognostic biomarker for drug adaptation.

Dual specific phosphatases (DUSPs) act downstream of BRAF on phosphorylated MAPK members to provide attenuation of signal. Loss of DUSP activity results in constitutive activation of the pathway. Prominent members of this family DUSP1 and DUSP2 are consistently downregulated at the transcriptional level. In prior clinical studies, somatic mutation of DUSP4 in MAPKiR has been reported [37]. Although in that case a genomic mechanism of resistance has been utilized, the outcome of reduced DUSP activity by genomic or transcriptomic changes is the same and leads to persistent triggering of MAPK effectors.

Metabolic genes support the rewiring of acquired resistance and have been shown to play an intricate role in the malignancy of skin cutaneous tissues. Glutamine and glucose metabolism showed sensitivity to combinations of MAPKi and metabolic inhibitors in preclinical studies [41]. The transciptomic profiles identified key enzymes in related, branching glycolytic pathways of serine, folate and pyrimidine metabolism. A cancer systems biology analysis of skin cutaneous melanoma brought a new master regulator and diagnostic target in cancer metabolism forward. Somatic mutations of DPYD have the ability to reconfigure and activate pyrimidine metabolism promoting rapid cellular proliferation and metastatic progression [42].

The forkhead box family of transcription factors is an important downstream target of the MAPK pathway and is currently being considered as a new therapeutic target in cancer, including melanoma therapy [43]. In epithelial cells, these transcriptional factors are directly involved in the expression of cyclin dependent kinase inhibitors and CDKN2A gene under the control of TGFβ [44, 45]. Both downregulation of anti-apoptotic targets as well as activation of proliferative metabolism have been observed as mechanisms contributing to MAPKiR. Downregulation of FOXF2 has been shown to promote cancer progression, EMT, and metastatic invasion [46]. In contrast, a different member of the FOX family, the stem cell transcription factor forkhead box D3 (FOXD3) has been identified as an adaptive mediator of the response to MAPK pathway inhibition selectively in mutant BRAF melanomas [47, 48].

To this point, we have identified non-genomic rewiring of pathways by RNA-Seq data and validated gene candidates in two cell lines by transcriptomics arrays. However, eventually perturbation of the identified resistance pathways by drug molecules or small hairpin RNAs will be needed to solidify a translational impact of candidate genes. Nevertheless, the established cell culture models of treatment resistance provide a broadly applicable platform to utilize high-throughput screening tools in the search for effective combinations of targeted therapies in cancer.

## Conclusion

The MAPK pathway undergoes major rewiring at the transcriptional level while acquiring inhibitor resistance. The outcome of such transcriptional plasticity is dysregulation at the level of different upstream master regulators, while maintaining similar effector genes. Combination therapies including targeted approaches and immune checkpoint inhibition are promising and rapidly improving. For these therapies to show durable, progression-free successes in the clinical setting, adaptation mechanisms of treatment resistances need to be understood. Cellular model systems in combination with transcriptome-wide analyses provide insight into how non-genomic drug adaptation is accomplished. Ongoing efforts are focused on utilizing the established preclinical models to overcome drug adaptation as well as precision medicine profiling of cancer patients. Over time, a better understanding of mechanisms involved in drug adaptation is likely to improve the effectiveness of melanoma therapy by delaying or controlling acquired resistance.

## Availability of data and material

Supplementary tables 1-4 are compiled in the submitted Supplementary information.

## Competing interests

There is no competing financial interest. F. L. M. is co-Founder and Medical director of Cancer Prevention Pharmaceuticals with no direct implications on the conducted study on melanoma resistance.

## Funding

F.V.F. is grateful for the support of grant CA154887 from the National Institutes of Health, National Cancer Institute. The research of the University of California Merced Systems Biology and Cancer Metabolism Laboratory is generously supported by University of California, Cancer Research Coordinating Committee CRN-17-427258, National Science Foundation, University of California Senate Graduate Research Council, and Health Science Research Institute program grants. F. L.-S. is supported by grant CA160756 from the National Institutes of Health, National Cancer Institute. F. L. M. and F. L.-S. are in part supported by the Waltmar and Oxnard Foundations and Aldrich Chair Endowment.

## Acknowledgements

We would like to thank Angela Garcia, Charles Fagundes, Garja Suner, Sandeep Sanghera, Taran Kaur, Kirandeep Kaur, Keedrian Olmstead, Stephen Wilson, and Rohit Gupta for help with maintaining the cellular models of drug resistant cancer cells.

## References

1. Davies H, Bignell GR, Cox C, Stephens P, Edkins S, Clegg S et al. Mutations of the BRAF gene in human cancer. Nature. 2002;417(6892):949–54. doi:10.1038/nature00766

2. Pollock PM, Harper UL, Hansen KS, Yudt LM, Stark M, Robbins CM et al. High frequency of BRAF mutations in nevi. Nat Genet. 2003;33(1):19–20. doi:10.1038/ng1054

3. Guan J, Gupta R, Filipp FV. Cancer systems biology of TCGA SKCM: efficient detection of genomic drivers in melanoma. Sci Rep. 2015;5:7857. doi:10.1038/srep07857

4. Tsai J, Lee JT, Wang W, Zhang J, Cho H, Mamo S et al. Discovery of a selective inhibitor of oncogenic B-Raf kinase with potent antimelanoma activity. Proc Natl Acad Sci U S A. 2008;105(8):3041–6. doi:10.1073/pnas.0711741105

5. Chapman PB, Hauschild A, Robert C, Haanen JB, Ascierto P, Larkin J et al. Improved survival with vemurafenib in melanoma with BRAF V600E mutation. N Engl J Med. 2011;364(26):2507–16. doi:10.1056/NEJMoa1103782.

6. Flaherty KT, Robert C, Hersey P, Nathan P, Garbe C, Milhem M et al. Improved survival with MEK inhibition in BRAF-mutated melanoma. N Engl J Med. 2012;367(2):107–14. doi:10.1056/NEJMoa1203421.

7. Flaherty KT, Infante JR, Daud A, Gonzalez R, Kefford RF, Sosman J et al. Combined BRAF and MEK inhibition in melanoma with BRAF V600 mutations. N Engl J Med. 2012;367(18):1694–703. doi:10.1056/NEJMoa1210093.

8. Robert C, Karaszewska B, Schachter J, Rutkowski P, Mackiewicz A, Stroiakovski D et al. Improved overall survival in melanoma with combined dabrafenib and trametinib. N Engl J Med. 2015;372(1):30–9. doi:10.1056/NEJMoa1412690.

9. Long GV, Stroyakovskiy D, Gogas H, Levchenko E, de Braud F, Larkin J et al. Combined BRAF and MEK inhibition versus BRAF inhibition alone in melanoma. N Engl J Med. 2014;371(20):1877–88. doi:10.1056/NEJMoa1406037.

10. Cheng Y, Zhang G, Li G. Targeting MAPK pathway in melanoma therapy. Cancer Metastasis Rev. 2013;32(3-4):567–84. doi:10.1007/s10555-013-9433-9.

11. Nazarian R, Shi H, Wang Q, Kong X, Koya RC, Lee H et al. Melanomas acquire resistance to B-RAF(V600E) inhibition by RTK or N-RAS upregulation. Nature. 2010;468(7326):973–7. doi:10.1038/nature09626

12. Wagle N, Emery C, Berger MF, Davis MJ, Sawyer A, Pochanard P et al. Dissecting therapeutic resistance to RAF inhibition in melanoma by tumor genomic profiling. J Clin Oncol. 2011;29(22):3085–96. doi:10.1200/JCO.2010.33.2312

13. Poulikakos PI, Persaud Y, Janakiraman M, Kong X, Ng C, Moriceau G et al. RAF inhibitor resistance is mediated by dimerization of aberrantly spliced BRAF(V600E). Nature. 2011;480(7377):387–90. doi:10.1038/nature10662

14. Van Allen EM, Wagle N, Sucker A, Treacy DJ, Johannessen CM, Goetz EM et al. The genetic landscape of clinical resistance to RAF inhibition in metastatic melanoma. Cancer Discov. 2014;4(1):94–109. doi:10.1158/2159-8290.CD-13-0617

15. Moriceau G, Hugo W, Hong A, Shi H, Kong X, Yu CC et al. Tunable-combinatorial mechanisms of acquired resistance limit the efficacy of BRAF/MEK cotargeting but result in melanoma drug addiction. Cancer Cell. 2015;27(2):240–56. doi:10.1016/j.ccell.2014.11.018

16. Hugo W, Shi H, Sun L, Piva M, Song C, Kong X et al. Non-genomic and Immune Evolution of Melanoma Acquiring MAPKi Resistance. Cell. 2015;162(6):1271–85. doi:10.1016/j.cell.2015.07.061

17. Emmons MF, Faiao-Flores F, Smalley KS. The role of phenotypic plasticity in the escape of cancer cells from targeted therapy. Biochem Pharmacol. 2016;122:1–9. doi:S0006-2952(16)30140-X [pii]

18. Lito P, Rosen N, Solit DB. Tumor adaptation and resistance to RAF inhibitors. Nat Med. 2013;19(11):1401–9. doi:10.1038/nm.3392

19. Carey TE, Takahashi T, Resnick LA, Oettgen HF, Old LJ. Cell surface antigens of human malignant melanoma: mixed hemadsorption assays for humoral immunity to cultured autologous melanoma cells. Proc Natl Acad Sci U S A. 1976;73(9):3278–82.

20. Fogh J, Fogh JM, Orfeo T. One hundred and twenty-seven cultured human tumor cell lines producing tumors in nude mice. J Natl Cancer Inst. 1977;59(1):221–6.

21. Giard DJ, Aaronson SA, Todaro GJ, Arnstein P, Kersey JH, Dosik H et al. In vitro cultivation of human tumors: establishment of cell lines derived from a series of solid tumors. J Natl Cancer Inst. 1973;51(5):1417–23.

22. Xing F, Persaud Y, Pratilas CA, Taylor BS, Janakiraman M, She QB et al. Concurrent loss of the PTEN and RB1 tumor suppressors attenuates RAF dependence in melanomas harboring (V600E)BRAF. Oncogene. 2012;31(4):446–57. doi:10.1038/onc.2011.250

23. Langmead B, Salzberg SL. Fast gapped-read alignment with Bowtie 2. Nat Methods. 2012;9(4):357–9. doi:10.1038/nmeth.1923

24. Robinson MD, McCarthy DJ, Smyth GK. edgeR: a Bioconductor package for differential expression analysis of digital gene expression data. Bioinformatics. 2010;26(1):139–40. doi:10.1093/bioinformatics/btp616

25. Mootha VK, Lindgren CM, Eriksson KF, Subramanian A, Sihag S, Lehar J et al. PGC-1alpha-responsive genes involved in oxidative phosphorylation are coordinately downregulated in human diabetes. Nat Genet. 2003;34(3):267–73. doi:10.1038/ng1180

26. Subramanian A, Tamayo P, Mootha VK, Mukherjee S, Ebert BL, Gillette MA et al. Gene set enrichment analysis: a knowledge-based approach for interpreting genome-wide expression profiles. Proc Natl Acad Sci U S A. 2005;102(43):15545–50. doi:0506580102 [pii]

27. Friedmann PS, Gilchrest BA. Ultraviolet radiation directly induces pigment production by cultured human melanocytes. J Cell Physiol. 1987;133(1):88–94. doi:10.1002/jcp.1041330111.

28. Liu F, Singh A, Yang Z, Garcia A, Kong Y, Meyskens FL, Jr. MiTF links Erk1/2 kinase and p21 CIP1/WAF1 activation after UVC radiation in normal human melanocytes and melanoma cells. Mol Cancer. 2010;9:214. doi:10.1186/1476-4598-9-214

29. Bradford MM. A rapid and sensitive method for the quantitation of microgram quantities of protein utilizing the principle of protein-dye binding. Anal Biochem. 1976;72:248–54. doi:S0003269776699996 [pii].

30. Sondergaard JN, Nazarian R, Wang Q, Guo D, Hsueh T, Mok S et al. Differential sensitivity of melanoma cell lines with BRAFV600E mutation to the specific Raf inhibitor PLX4032. J Transl Med. 2010;8:39. doi:10.1186/1479-5876-8-39

31. Shain AH, Bastian BC. From melanocytes to melanomas. Nat Rev Cancer. 2016;16(6):345–58. doi:10.1038/nrc.2016.37

32. Tiffen J, Wilson S, Gallagher SJ, Hersey P, Filipp FV. Somatic Copy Number Amplification and Hyperactivating Somatic Mutations of EZH2 Correlate With DNA Methylation and Drive Epigenetic Silencing of Genes Involved in Tumor Suppression and Immune Responses in Melanoma. Neoplasia. 2016;18(2):121–32. doi:10.1016/j.neo.2016.01.003

33. Lito P, Pratilas CA, Joseph EW, Tadi M, Halilovic E, Zubrowski M et al. Relief of profound feedback inhibition of mitogenic signaling by RAF inhibitors attenuates their activity in BRAFV600E melanomas. Cancer Cell. 2012;22(5):668–82. doi:10.1016/j.ccr.2012.10.009

34. Yao Z, Torres NM, Tao A, Gao Y, Luo L, Li Q et al. BRAF Mutants Evade ERK-Dependent Feedback by Different Mechanisms that Determine Their Sensitivity to Pharmacologic Inhibition. Cancer Cell. 2015;28(3):370–83. doi:10.1016/j.ccell.2015.08.001

35. Wagle N, Van Allen EM, Treacy DJ, Frederick DT, Cooper ZA, Taylor-Weiner A et al. MAP kinase pathway alterations in BRAF-mutant melanoma patients with acquired resistance to combined RAF/MEK inhibition. Cancer Discov. 2014;4(1):61–8. doi:10.1158/2159-8290.CD-13-0631

36. Shi H, Hugo W, Kong X, Hong A, Koya RC, Moriceau G et al. Acquired resistance and clonal evolution in melanoma during BRAF inhibitor therapy. Cancer Discov. 2014;4(1):80–93. doi:10.1158/2159-8290.CD-13-0642

37. Johnson DB, Menzies AM, Zimmer L, Eroglu Z, Ye F, Zhao S et al. Acquired BRAF inhibitor resistance: A multicenter meta-analysis of the spectrum and frequencies, clinical behaviour, and phenotypic associations of resistance mechanisms. Eur J Cancer. 2015;51(18):2792–9. doi:10.1016/j.ejca.2015.08.022

38. Filipp FV. Precision medicine driven by cancer systems biology. Cancer Metastasis Rev. 2017;36(1):1–18. doi:10.1007/s10555-017-9662-4.

39. Johannessen CM, Johnson LA, Piccioni F, Townes A, Frederick DT, Donahue MK et al. A melanocyte lineage program confers resistance to MAP kinase pathway inhibition. Nature. 2013;504(7478):138–42. doi:10.1038/nature12688

40. Garraway LA, Widlund HR, Rubin MA, Getz G, Berger AJ, Ramaswamy S et al. Integrative genomic analyses identify MITF as a lineage survival oncogene amplified in malignant melanoma. Nature. 2005;436(7047):117–22. doi:nature03664 [pii]

41. Hernandez-Davies JE, Tran TQ, Reid MA, Rosales KR, Lowman XH, Pan M et al. Vemurafenib resistance reprograms melanoma cells towards glutamine dependence. J Transl Med. 2015;13:210. doi:10.1186/s12967-015-0581-2

42. Edwards L, Gupta R, Filipp FV. Hypermutation of DPYD Deregulates Pyrimidine Metabolism and Promotes Malignant Progression. Mol Cancer Res. 2016;14(2):196–206. doi:10.1158/1541-7786.MCR-15-0403

43. Yang JY, Hung MC. A new fork for clinical application: targeting forkhead transcription factors in cancer. Clin Cancer Res. 2009;15(3):752–7. doi:10.1158/1078-0432.CCR-08-0124

44. Gomis RR, Alarcon C, He W, Wang Q, Seoane J, Lash A et al. A FoxO-Smad synexpression group in human keratinocytes. Proc Natl Acad Sci U S A. 2006;103(34):12747–52. doi:0605333103 [pii]

45. Lasfar A, Cohen-Solal KA. Resistance to transforming growth factor beta-mediated tumor suppression in melanoma: are multiple mechanisms in place? Carcinogenesis. 2010;31(10):1710–7. doi:10.1093/carcin/bgq155

46. Kong PZ, Li GM, Tian Y, Song B, Shi R. Decreased expression of FOXF2 as new predictor of poor prognosis in stage I non-small cell lung cancer. Oncotarget. 2016;7(34):55601–10. doi:10.18632/oncotarget.10876

47. Abel EV, Aplin AE. FOXD3 is a mutant B-RAF-regulated inhibitor of G(1)-S progression in melanoma cells. Cancer Res. 2010;70(7):2891–900. doi:10.1158/0008-5472.CAN-09-3139

48. Abel EV, Basile KJ, Kugel CH, 3rd, Witkiewicz AK, Le K, Amaravadi RK et al. Melanoma adapts to RAF/MEK inhibitors through FOXD3-mediated upregulation of ERBB3. J Clin Invest. 2013;123(5):2155–68. doi:10.1172/JCI65780

